# Simultaneous 3D super-resolution fluorescence microscopy and atomic force microscopy: combined SIM and AFM platform for cell imaging

**DOI:** 10.1101/638262

**Authors:** Ana I. Gómez-Varela, Dimitar R. Stamov, Adelaide Miranda, Rosana Alves, Cláudia Barata-Antunes, Daphné Dambournet, David G. Drubin, Sandra Paiva, Pieter A. A. De Beule

## Abstract

Correlating data from different microscopy techniques holds the potential to discover new facets of signaling events in cellular biology. Here we report for the first time a hardware set-up capable of achieving simultaneous imaging of spatially correlated super-resolution fluorescence microscopy and atomic force microscopy, a feat only obtained until now by fluorescence microscopy set-ups with spatial resolution restricted to the Abbe resolution limit. We hereby remove the need to perform independent measurement and subsequent data averaging required to eliminate cell-to-cell variation in observed signals. We detail system integration, demonstrate system performance and report imaging of sub-resolution fluorescent beads and genome-engineered human bone osteosarcoma epithelial cells.

## 1. Introduction

Visualization of biological specimens can be accomplished through several available microscopy techniques, each of them presenting inherent benefits and constraints. The combination of different imaging platforms can further extend their applications [1, 2]. The major significance of correlative imaging relies on its capacity to provide complementary information such as morphological, chemical and biophysical details from the same sample during a single experiment.

Atomic Force Microscope (AFM) was introduced in 1986 by Binning *et al.* [3], being a high-resolution surface probe technique that allows topographical and nano-mechanical characterisation. AFM has become a powerful tool to team up with optical microscopes, especially regarding the biosciences field. On the other hand, super-resolution fluorescence optical microscopy methods have been a major breakthrough for biological samples examination, having being actively developed in the latter years to surpass the optical diffraction barrier [4]. Nonetheless, they are limited in some aspects such as for example nanomechanical sample characterization. Therefore, a correlative microscopy approach is bene cial as it allows to pair up the high spatial resolution and mechanical characterization potential of an AFM, with the additional molecular information available, through various fluorescent markers or immunostains [5, 6].

Despite the great potential on simultaneous operation of AFM and super-resolution fluorescence microscopy, no one to the best of our knowledge has reported it. Current AFM techniques combining optical fluorescence imaging normally perform successive measurements and then superimpose the images obtained from the separate acquisitions. Main reasons for this are: i) imaging in a simultaneous mode causes perturbation on the AFM cantilever operation due to fluorescence excitation light, and ii) noise transfer from the optical microscope can disturb AFM measurements [7, 8].

The atomic force microscope has been successfully integrated with different optical microscopy techniques [9, 10], allowing to overcome some of its individual limitations when it comes to, for instance, sample penetration and biological specificity. In this regard, AFM has been combined with Confocal Laser Scanning Microscopy (CLSM) [11], Aperture Correlation Microscopy [8], Total Internal Reflection Fluorescence Microscopy (TIRFM) [12], and Fluorescence Lifetime Imaging (FLIM) [13]. AFM has also been integrated with super-resolution microscopy schemes, namely Stimulated Emission Depletion (STED) [14, 15], Photoactivated Localization Microscopy (PALM)[16] and Stochastic Optical Reconstruction Microscopy (STORM) [17, 18]. The integration of AFM with fluorescence super-resolution schemes faces a variety of problems impeding simultaneous data acquisition. For instance, high-powered light sources interact strongly with gold coated AFM cantilevers, causing excessive heating or even cantilever coating deterioration. High powered light sources are commonplace for the depletion beam of a STED set-up as well as accelerators for a fast read-out in STORM microscopy. Furthermore, STORM microscopy requires uorophores exhibiting blinking behaviour which is typically promoted by immersing the sample in a buffer containing an enzymatic oxygen scavenger. The latter precludes simultaneous operation of the AFM and fluorescent measurements, as the ingredients of the buffer stick to the AFM cantilever, so correlative imaging is normally performed by acquiring first the AFM image and then adding the buffer for super-resolution microscopy [16, 19]. Recent research has proposed a new dye to circumvent this problem that allows correlative AFM and STORM imaging without the need to change the buffer [18].

Here, we report on a new super-resolution microscopy platform that allows to perform simultaneously fluorescence microscopy and atomic force microscopy. In this context, we combine AFM with Structured Illumination Microscopy (SIM), a super-resolution technique that allows to surpass the diffraction barrier by a factor of two in every spatial direction [20]. Contrary to other super-resolution modalities, SIM relies on a fluorescent excitation light pattern favourable to avoid AFM cantilever disruption during simultaneous operation, as we reported previously on a novel system for synchronic fluorescence optical sectioning microscopy through aperture correlation microscopy using a differential Spinning Disk (DSD) and AFM [8].

The principle of SIM relies on the projection of a grating onto the sample and the subsequent reconstruction of the high-resolution image from different grating imaging positions. This technique has been used for biological imaging, in particular for fixed samples, but it has also proven to be easily applicable to live cell imaging [21, 22]. SIM is a very promising imaging modality to reveal dynamic processes in live biological samples in 3D and presents some advantages over other super-resolution techniques as it does not require the use of specialized uorophores or sample preparations, induces less phototoxicity and can reach higher acquisition rates for longer periods of time as compared to PALM/STORM and STED [23]. Because SIM requires a relatively low illumination power, its integration with AFM is very appropriate for non-disruptive simultaneous operation. Aside, as SIM is subject to su er from artifacts during image reconstruction process [24, 25], a combination with AFM can help to validate super-resolution data.

To demonstrate system performance, we first image a sample consisting of sub-resolution fluorescent beads. SIM-AFM simultaneous operation is then explored using CRISPR/Cas9 genome-edited human cells expressing a fluorescently tagged plasma membrane transporter from their native genomic loci. This model is particularly relevant for super-resolution microscopy techniques as it avoids the typical overexpression-induced artefacts that often affects protein localization and dynamics [23].

## 2. Materials and Methods

### 2.1. SIM-AFM set-up

The combined SIM-AFM platform is based on an atomic force microscope (JPK NanoWizard 3, Bruker Nano GmbH, Berlin, Germany) and a structured illumination microscope (N-SIM E, Nikon Instruments Europe B.V., Amsterdam, The Netherlands). The schematic set-up of the system is shown in Figure 1. The AFM is mounted on an inverted microscope (Eclipse Ti2-E, Nikon Instruments Europe B.V., Amsterdam, The Netherlands) and standard monochrome CCD camera (ProgRes^®^ MFCool, Jenoptik, Jena, Germany) is coupled for AFM laser spot cantilever alignment. An infrared laser filter is mounted on the binocular phototube of the Ti2 body for maintaining ocular safety and an extra long working distance (ELWD) lens of 75 mm (T1-CELWD ELWD, Nikon Instruments Europe B.V., Amsterdam, The Netherlands) is installed in the system.

**Figure 1:**
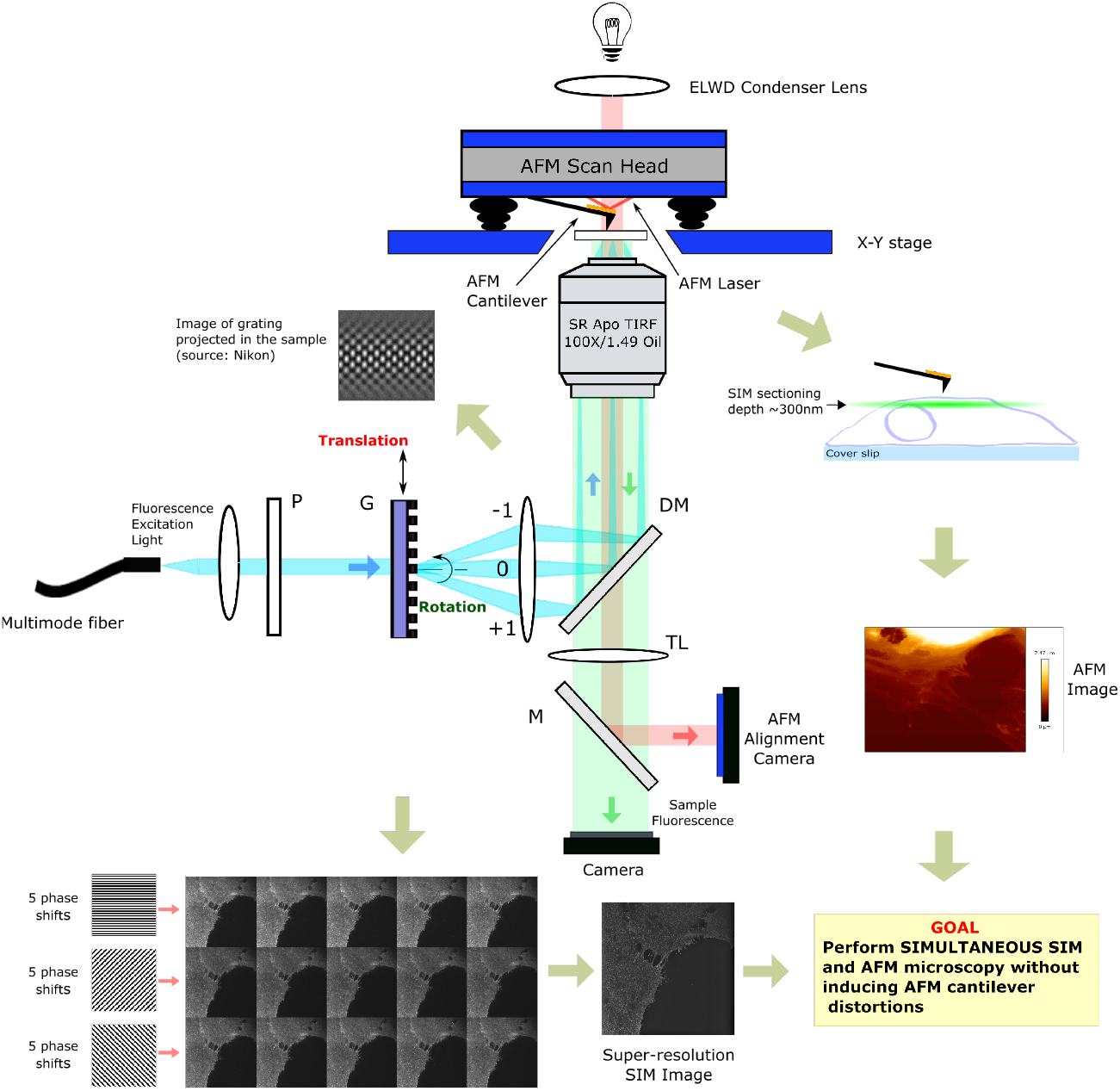
Simplified schematic set-up of the SIM-AFM system. In this case, structured illumination is generated using a grating (G). Collimated and expanded laser light illuminates the grating, resulting in the diffraction of multiple orders. Only 0^th^ and ±1^st^ orders are allowed into the illumination path and focused on the back focal plane of the objective and the created stripped illumination pattern excites the sample. Collection of the fluorescence signal is achieved by a high aperture microscope objective, a dichromatic mirror (DM) and a tube lens (TL). The AFM laser is reflected by a mirror (M) to a high-dynamic range camera coupled to a microscope port for cantilever laser spot alignment. At the same time, nanomechanical mapping of the sample is performed by a probe consisting of a flexible cantilever and a tip. After simultaneous SIM and AFM images acquisition, they are combined into a single image.

The AFM was operated in an advanced force-spectroscopy based mode, referred to as Quantitative Imaging (QI ™), which allows nanomechanical characterisation and simultaneous imaging [26]. For static measurements on beads we used force modulation cantilevers (FM, NanoWorld, Switzerland), with a nominal resonance frequency of 75 kHz in air, spring constant of 2.8 N m^−1^, reflective detector gold coating, and monolithic silicon pyramidal tips with radius of curvature (ROC) of 8 nm. For measurements on cells in liquid, we used qp-BioAC-CI-CB1 cantilevers (NanoSensors, Switzerland), with a nominal resonance frequency of 90 kHz (in air), spring constant of 0.3 N m^−1^, partial gold coating on the detector side, and quartz-like circular symmetric hyperbolic (double-concaved) tips with ROC of 30 nm.

SIM illumination is provided by laser light previously coupled into a multimodal fiber. In the Nikon N-SIM E microscope three different wavelengths are available (488/561/640nm). The output collimated light from the fiber travels towards a grating. Only the zero, first positive, and first negative order diffracted light (3D SIM mode) are allowed to pass by a blocking element, resulting in a grating projected onto the sample that fades away into a blur a short distance away from focus, favourable for providing constant illumination of the AFM cantilever. For fluorescence light collection, a high aperture microscope objective (CFI SR APO TIRF 100× Oil N.A. 1.49, WD 0.12 mm, Nikon Instruments Europe B.V., Amsterdam, The Netherlands) is used. Super-resolution information is recorded with a scientific CMOS (sCMOS) camera (C11440-42U Orca Flash4.0 LT, Hamamatsu Photonics K.K., Boston, MA). The final image is reconstructed by computational methods (NIS-Elements, Nikon Instruments Europe B.V., Amsterdam, The Netherlands). The reconstruction process uses 15 raw images obtained at different orientations of the structured illumination, which is done by moving the diffraction grating (5 phase shifts and 3 rotations around the optical axis). All elements are controlled with NIS Elements AR from Nikon Instruments.

The entire platform (Figure 2) is mounted on an optical table with active isolator leg bundles (T1225Q - Nexus Optical Table, Thorlabs, Inc., Newton, NJ, USA). All experiments were carried out at 21 °C.

**Figure 2:**
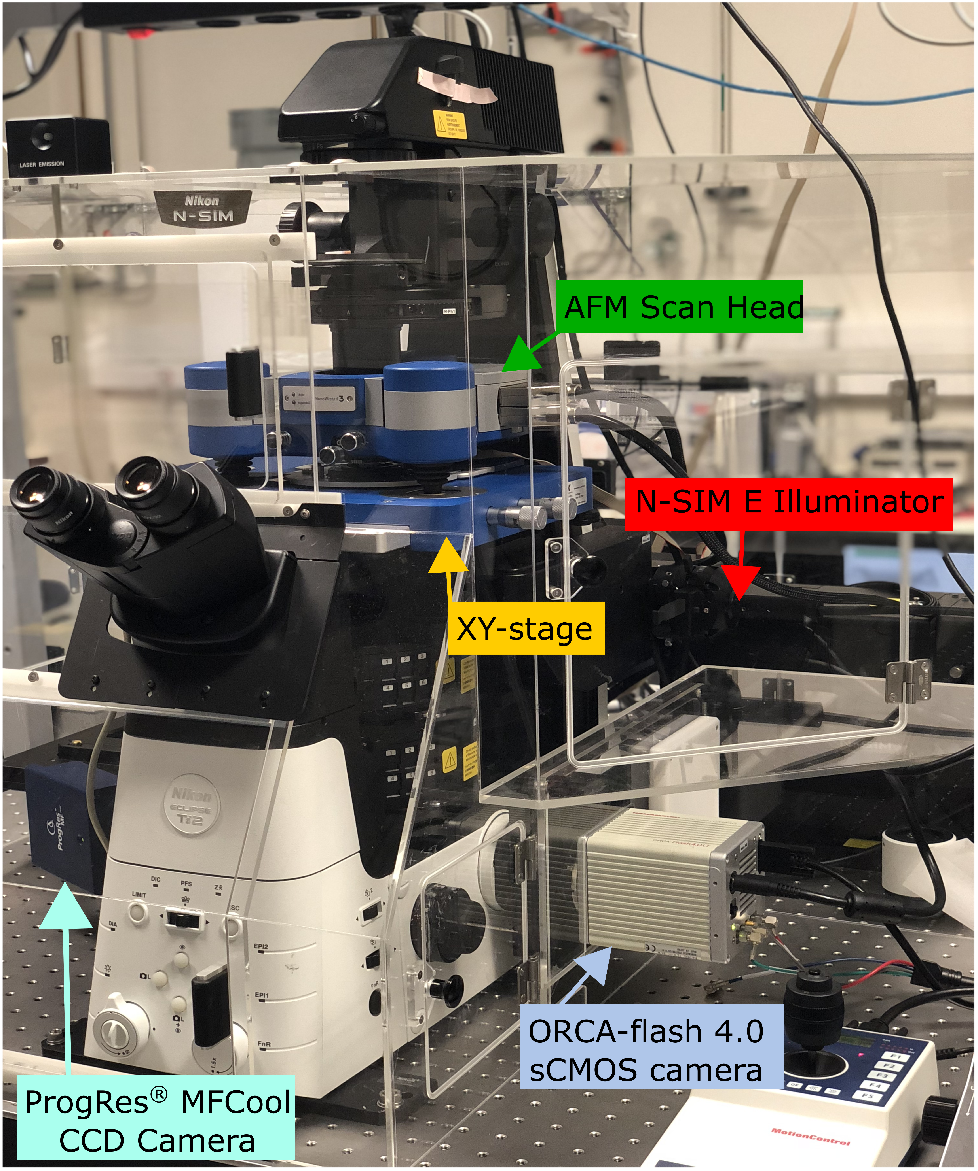
Image of the SIM-AFM system. The N-SIM illuminator module provides the structured illumination. AFM scan head (XYZ) and sample positioning stage (XY) are mounted on the inverted optical microscope. Laser spot alignment of the AFM is accomplished using a standard monochrome CCD camera while fluorescence signal is acquired with a scientific CMOS camera.

### 2.2. Sample preparation

#### 2.2.1. Fluorescent beads

Commercially available FluoSpheres^®^ beads (carboxylate-modified orange fluospheres, 0.1 μm) from Thermo Fisher Scientific were used as test samples. A stock suspension containing 0.5 μL fluorescent beads in 25 μL of ethanol was prepared and then vortexed for 5 min. The stock solution was further diluted to 1:100 in Milli-Q water and 500 μL of the beads suspension was deposited on fluorodish™ glass dish (bottom poly-D-Lysine coated with a glass thickness of 0.17 mm) and left to air dry at room temperature. The glass bottom dish was thoroughly cleaned on the side of the objective immersion medium with ethanol prior to measurements.

#### 2.2.2. Cell culture and genome editing

In this work, genome-engineered cancer cells were applied to demonstrate the SIM-AFM simultaneous operation. Gene-edited U2OS cells expressing Monocarboxylate Transporter 1 fluorescently labelled with Enhanced Green Fluorescent Protein (MCT1-EGFP) in both alleles were grown in DMEM/F12/GlutaMAX (Cat. No. 10565042, Gibco, Waltham, MA) supplemented with 10% fetal bovine serum (Cat. No. 35-010-CV, Corning) and Penicillin-Streptomycin 100× solution (Cat. No. 15140-122, Gibco, Waltham, MA). 24 h before imaging, the cells were seeded on a *μ*-Dish 35 mm, high ibiTreat (Ibidi, Martinsried, Germany). Cells were then fixed in 4 % paraformaldehyde at room temperature for 20 min and washed with Dulbecco’s phosphate-buffered saline (Cat. No. 14190-144, Gibco, Waltham, MA).

The MCT1 gene was targeted at the N-terminal using CRISPR/Cas9 genome editing technology and 5’-GTAGATAAATTCCAAAATGC-3’ as a guide RNA, both expressed utilizing pX330 plasmid [27, 28]. 2 × 10^6^ U2OS cells were electroporated with 5 μg of pX330 Cas9 plasmid and 15 μg of donor plasmid. Cells were sorted out by an Inux sorter (BD Bioscience) for GFP fuorescence 72 h after electroporation. Clonal populations were isolated and characterized, as previously described [29]. Sequencing revealed a one nucleotide sequence variation, which does not alter the protein or intron sequences.

### 2.3. Image registration

The simultaneous operation of AFM with optical microscopy enables the collection of optical sectioning fluorescence and nano-mechanical mapping information from a sample. However, one complication of SIM is that, in order to avoid artifacts in the final image, requires of optimized experimental implementation, consideration of bleaching properties of the sample [30, 31] and proper selection of reconstruction parameters [32, 33], as SIM has a need of a complex post-processing step. Hence, coupling of AFM and SIM can also be a powerful tool to validate the results obtained with the latter.

The successful combination of AFM with optical images is not a straight-forward process. Lenses in microscopes exhibit optical aberrations that distort the image, while AFM registers “real-space” images using highly linear piezo-electric elements [34]. To circumvent this problem and correctly ovarlay SIM data onto AFM images we use a software module (DirectOverlay™, JPK BioAFM, Bruker Nano GmbH, Berlin, Germany) to calibrate the optical image. In this calibration process the cantilever is displaced to a set of predefined coordinates in a 3 × 3 or a 5 × 5 grid pattern, registers an optical image at each position and a transform function between the AFM and the optical image is determined [8].

## 3. Results and Discussion

At first, we evaluated the spatial resolution improvement of the structured illumination system (Figure 3). To this end, we used both fluorescent beads and fixed cell samples. Both wide field (WF) and SIM images were acquired with a pixel size of 65 nm correspondingly at the sample plane, at starting resolution of 1024 × 1024 pixels. The positions at which arbritary cross-sections are drawn are marked in the WF panels (Figure 3a,d). Prior to analysis, all TIFF les underwent a 16-to-8bit conversion. In addition, due to the image reconstruction algorithm used, all 3D SIM images had doubled pixel resolution (2048 × 2048), which had to be resized and downscaled to the original WF pixel resolution by a factor of 2.

**Figure 3:**
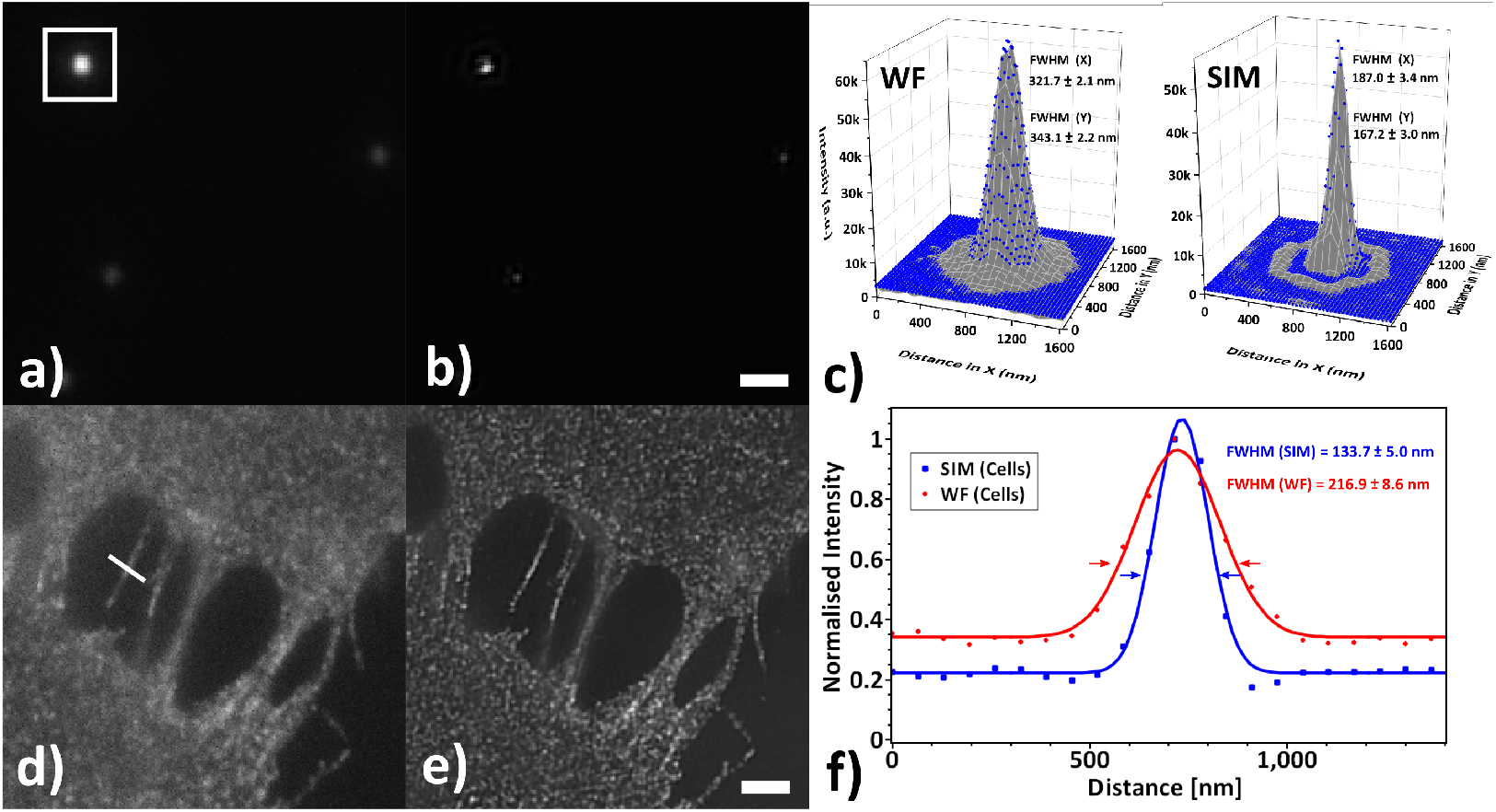
Improvement in optical resolution. (a) Wide field (WF) and (b) structured illumination microscopy (SIM) reconstructed images of orange fluorescent beads, 100 nm in diameter. (d) WF and (e) SIM images from gene-edited U2OS cells, labelled with EGFP. The intensity of the outlined regions in (a,b) were subjected to a non-linear surface tting with a Gaussian 2D function, allowing to compare the 3D PSFs in (c). Normalised intensity profiles from the arbitrary cross-sections in (d,e) were subsequently fit with a 1D Gaussian function to allow direct correlation of the FWHM for each sample in (f). Scalebars in (a,b) and (d,e) correspond to 1 μm and 2 μm respectively.

The outlined areas in Figure 3a,b were subjected to a surface fit with a Gaussian2D function (scaled Levenberg-Marquardt algorithm, tolerance of 10^−9^, resulting *R*^2^ of 0.91 for 3D SIM and 0.99 for WF respectively) in Origin v9.6.0.172 (OriginLab Corporation, Northampton, MA), allowing for an elliptical point spread function (PSF) [35]. Following a normalisation of the image intensity values in Figure 3d,e, the cross-section profiles were fit with a Gaussian function (scaled Levenberg-Marquardt algorithm, tolerance of 10^−4^, resulting *R*^2^ of 0.99) in SciDAVis v1.22 to determine the full width at half maximum (FWHM) of the corresponding PSF. For beads we observed a reduction in the averaged FWHM of the e ective focal spot from 332.4 nm to 177.1 nm, whereas for cells the FWHM was reduced from 216.9 nm to 133.7 nm, corresponding to a 1.88× and 1.62× increase in lateral resolution, approaching the theoretical maximum.

Once it was verified that it is possible to work below the diffraction limit, we proceeded with correlation of the optical and AFM data. The accuracy of the optical overlay relies on the XY-linearised closed loop positioning of the AFM piezos. By moving the tip to a set of fixed coordinates (3 × 3 or 5 × 5) in the AFM, it was possible to detect the tip position in the optical images and calculate a transfer function that allows for compensation of the non-linearities, and abberations in the optical images. Following that, it was possible to directly choose AFM regions of interest (ROI) from the calibrated optical images and collect data.

The correlation of the SIM channels and topographical AFM data is given in Figure 4. The overlay of the 100 nm beads with the SIM data, shown as semi-transparrent images, shows that the topographical centers of the beads overlap very well the intensity maxima of the bead positions from the SIM images. Evaluating the overlay accuracy on samples with defined morphology, such as beads or calibration standards, is preferable compared to large and structurally heterogeneous samples such as cells.

**Figure 4:**
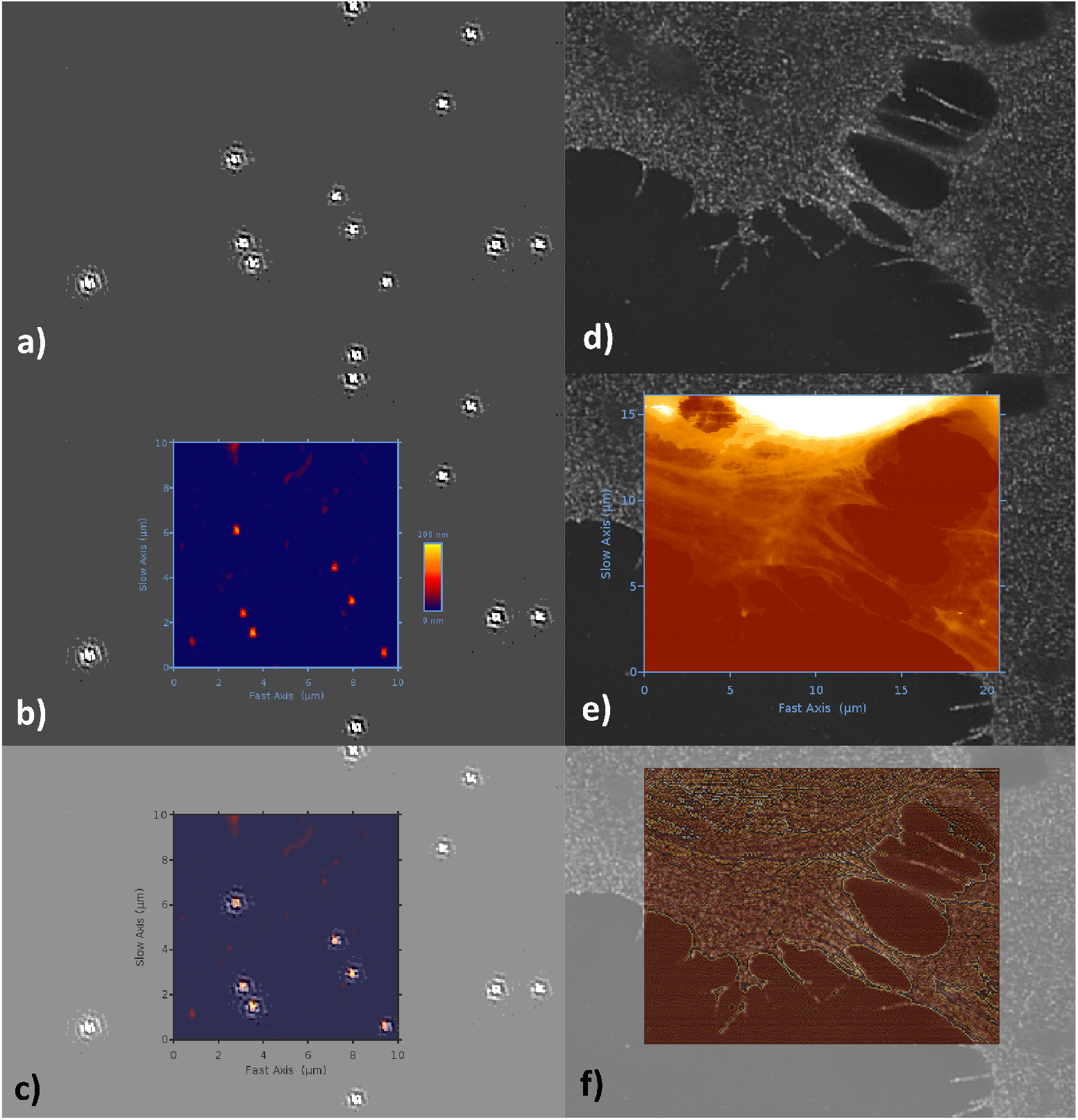
Correlation of the SIM and AFM data. The SIM images of the fluospheres (a) are superimposed with an AFM image (b) and subsequently correlated in a semi-transparent mode (c). Similarly, the optical data from the human bone osteosarcoma epithelial cells (d) is overlayed with the height/topography channel of the AFM data (e). The correlated SIM-AFM channels on cells (f), include a mean-curvature enhanced AFM-topographic information (see details in text).

Following the successful overlay on 100 nm beads, we proceeded with fixed human bone osteosarcoma cell line. Due to a significantly extended Z-scale of the AFM images (Figure 4e), it is not always straightforward to represent the morphological information from the leading cell surface edge features demonstrated here by the cytoskeletal framework, and the significantly higher nucleus in the upper half of the AFM images. In such cases, it is bene cial to represent the AFM images by using a pixel difference Z-scale filter of some sort. In this particular case, we have treated the AFM height image in Figure 4f with a curvature function [36] that highlights the curvature in the image by calculating the local mean curvature operation A for each pixel:

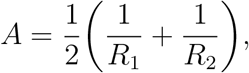

where *R*_1_ and *R*_2_ are the smallest and the largest radii determined at each pixel.

Correlating different types of datasets, such as optical and AFM data bodes a different set of questions and problems about the efficiency of multi-dimensional data representation. On one side, it is necessary to consider that both SIM and AFM techniques carry by de nition very different types of information, interpreted by the different operation principles and detection methods. This is further complicated by the different spatial and axial resolution of both techniques, which can differ in the range of up to 2 (spatial) or 3 (axial) orders of magnitude. Another practical limitation is the 3D-sectioning of the samples, as well as imaging penetration depth. Whereas in the case of SIM, 3D thinly sectioned images with axial resolution of down to 300 nm are now possible [37], the Z-scaling (not resolution) in conventional AFM images is typically con ned to the overall sample morphology. A certain solution to this problem is the application of force-spectroscopy based imaging (force-controlled AFM), which allows us to interpret the 3D-Force volume data of the acquired AFM dataset, i.e. each pixel has a force curve associated, and analytically subtract different quantitative information [38]. In this particular scenario, it is possible to determine the theoretical contact point in the individual force curve, and look at the sample topography or indentation information within the applied force range. This allows to look at the indentation phenomenon at a range of reference forces and enables a 3D tomographic/sectioning reconstruction of the sample within the studied indentation depth. We have given an example for that by sectioning part of the AFM image in Figure4e and representing the topography at different set of forces (Figure 5).

**Figure 5:**
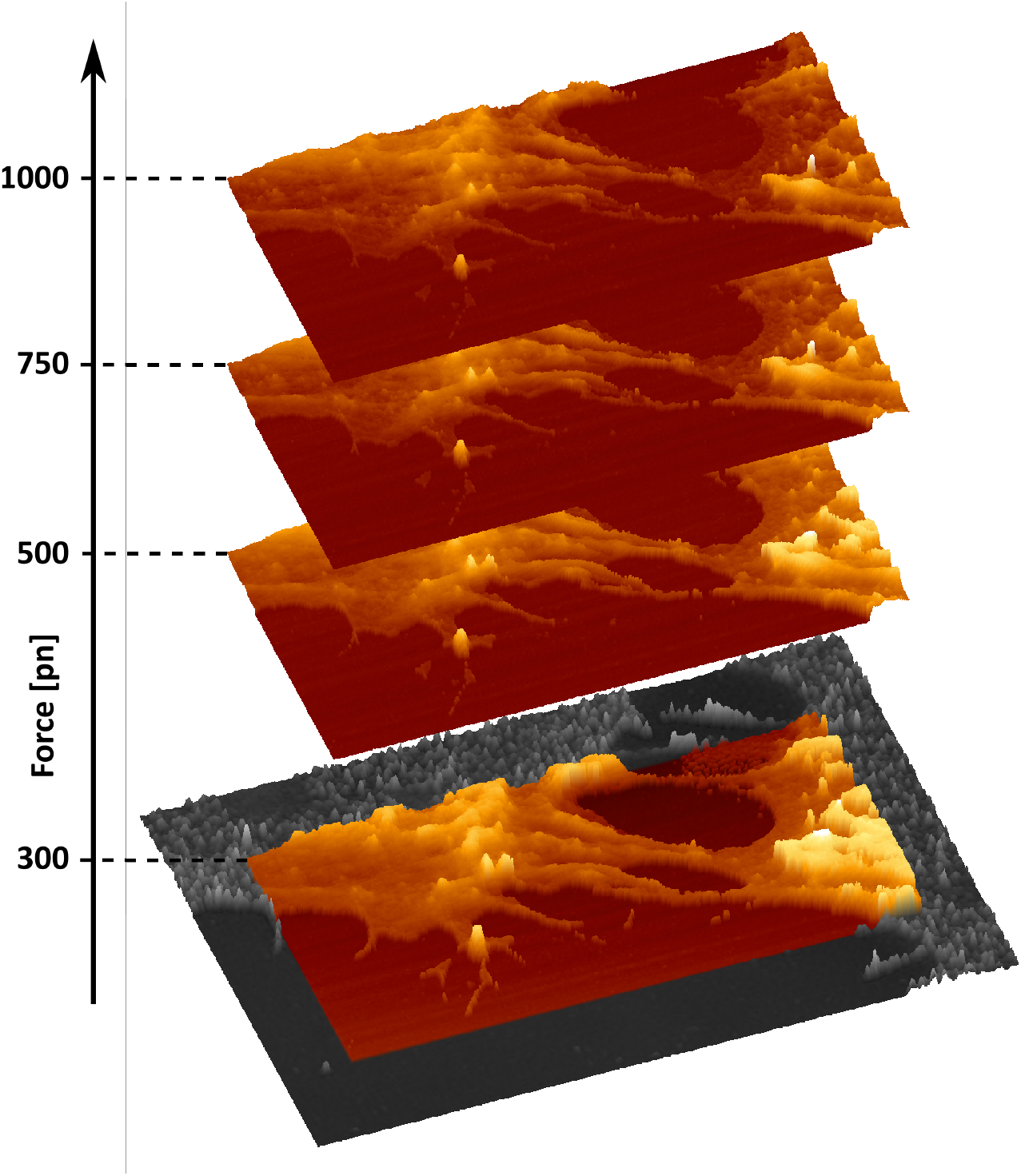
Multidimensional representation of correlated microscopy sets. Recording a complete set of force curves enables the calculation of the sample topography within the range of pre-defined force setpoint and reconstruct the surface structure of the cells. The overlay at 300 pN with the SIM channel is given for consideration of the spatial overlay accuracy. Size of the AFM image in X and Y is 20.74 μm and 10.37 μm respectively.

The tip scanning AFM design further enables a simultaneous acquisition of AFM and SIM images. We previously showed that for a simultaneous combination of an optical microscope equipped with a differential spinning disc and AFM, there are certain system-specific noise sources, which stem either from the AFM cantilever bimorph distortion induced by the different thermal expansion coe cients of cantilever material (*Si*/*Si*_3_*N*_4_) and reflective coatings (*Au*), or the pure mechanical noise transfer of the DSD system [8]. Our tests showed that there is no substantial difference within the noise response with the 3D N-SIM illumination in place (Figure 6). The results suggest that although the N-SIM illumination appears to have a small effect on the surface roughness parameters, it does not affect the acquisition or evaluation of individual structures on the cell surface. In addition, the used qp-bioAC-CI cantilevers have only a limited partial *Au* coating which considerably reduces the bending of the cantilevers during operation [8].

**Figure 6:**
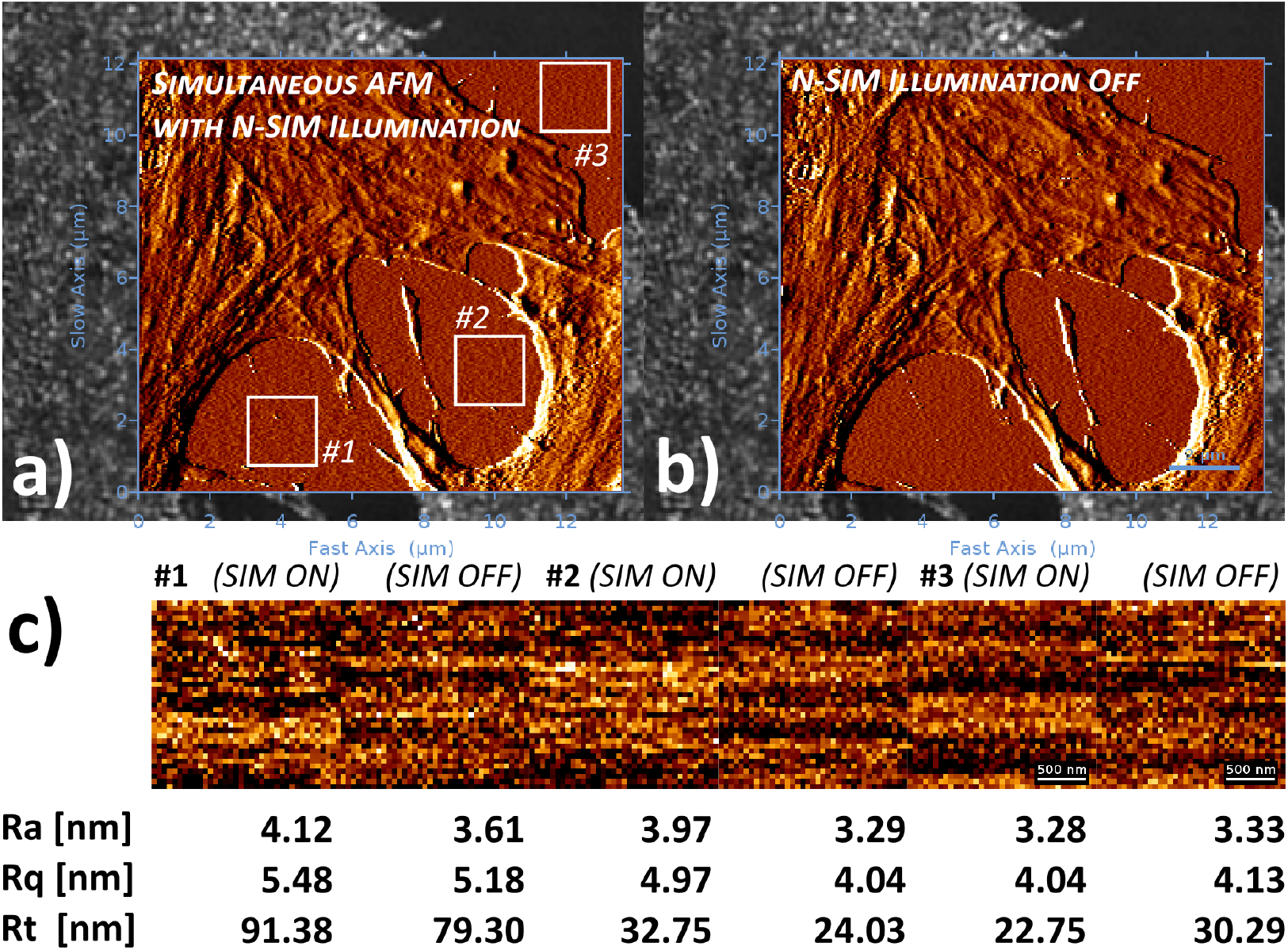
Simultaneous SIM/AFM acquisition. The AFM measurements were carried out on fixed USO2 cells in medium/buffer with (a) and without N-SIM illumination (b). For convenience and enhanced feature/noise contrast, both AFM topography images in the SIM-AFM overlays are displayed with an edge detection algorithm using a pixel difference operator in X. The topography images from petri dish surface on three positions (labelled in the gures) were planefit (1st order polynomial function) to compensate for tilts in the sample surface, and subjected to surface roughness analysis (c). For comparison reasons, the average roughness (Ra), RMS roughness (Rq) and peak-to-valley roughness (Rt) values are given below the corresponding height profiles.

## 4. Conclusion

A 3D SIM super-resolution system has been successfully integrated with a tip-scanning QI™ nanomechanical mechanical mapping AFM for simultaneous SIM-AFM operation. The combined SIM-AFM integration was tested on 100 nm fluorescent spheres and fixed osteosarcoma cells, demonstrating a good 2D spatial correlation of both signals. The spatial resolution obtained in both cases was approaching the theoretical SIM diffraction limit. We also give an example for multidimensional representation of correlated microscopy sets, by introducing section of the 3D AFM force-volume sets. Finally, the results obtained indicate that simultaneous SIM illumination during AFM cantilever operation does not distort the evaluation of the samples.

The simultaneous operation of AFM and super-resolution fluorescence microscopy technique provides a powerful observational tool on the nanoscale, albeit data acquisition is obstructed by a series of integration problems. We believe that the combination of SIM with AFM presents one of the most promising schemes enabling synchronous imaging, allowing the recording of nanomechanical data and cellular dynamics visualization at the same time.

## 5. Conflict of interest

There are no conflicts to declare.

## 6. Acknowledgements

Ana I. Gómez Varela wishes to acknowledge support from the Xunta de Galicia, Consellería de Cultura, Educación e Ordenación Universitaria e da Consellería de Economía, Emprego e Industria (Programa de axudas de apoio á etapa de formación posdoutoral 2017). Adelaide Miranda and Pieter De Beule acknowledge financial support from Norte’s Regional Operational Programme 2014-2020 Norte2020 (NORTE-01-0145-FEDER-000019). Sandra Paiva and Rosana Alves thank Fulbright Commission at Portugal and Luso-American Development Foundation for their financial support to perform research work at UC Berkeley, California, USA. We thank the U.S. Embassy in Portugal for supporting David Drubin’s visit to Portugal. We thank Ann Fisher of the UC Berkeley Cell Culture Facility for help with cell culture. We thank Dr. Kartoosh Heydari of the Cancer Research Lab Flow Cytometry Core Facility of UC Berkeley. Rosana Alves and Cláudia Barata-Antunes are recipients of PhD fellowships from the Portuguese Foundation for Science and Technology (PD/BD/113813/2015 and PD/BD/135208/2017, respectively).

The authors want to thank Nikon and Izasa Scientific for their support to the experiment by providing an E-SIM microscope set-up on loan. We also gratefully acknowledge Dr. Kees van der Oord from Nikon Instruments Europe B.V. for his assistance with the SIM microscope as well as Paulo Madureira and Carlos Pitães from IZASA Portugal and Jordi Recasens from IZASA Spain for their assistance with the integration of the SIM and AFM microscopes. Finally, we want to thank Benjamin Holmes for fruitful discussions and relentless support.

## Author contribution

Conception and design of the integration of the SIM and AFM microscopes was done by AIGV, AM and PDB. AIGV, DS, AM and PDB performed the hardware set-up of the microscopes. RA, CB, DD, DGR and SP were responsible of the genome-edited human cells. Experiments were conducted by AIGV, AM, DS and PDB. AIGV and DS wrote the manuscript with significant contributions from all authors. All authors gave final approval for publication.

## References

[1] C. Smith, Two microscopes are better than one, Nature 492 (2012) 293.

[2] J. Caplan, M. Niethammer, R. M. Taylor, K. J. Czymmek, The power of correlative microscopy: multi-modal, multi-scale, multi-dimensional, Current Opinion in Structural Biology 21 (2011) 686 – 693.

[3] G. Binnig, C. F. Quate, C. Gerber, Atomic force microscope, Physical Review Letters 56 (1986) 930–933.

[4] U. Endesfelder, Super-resolution microscopy. A practical guide. Von Udo J. Birk., Angewandte Chemie International Edition 57 (2018) 7939–7939.

[5] P. Bondia, S. Casado, C. Flors, Correlative super-resolution fluorescence imaging and atomic force microscopy for the characterization of biological samples, Springer New York, New York, NY, pp. 105–113.

[6] L. A. Chtcheglova, P. Hinterdorfer, Simultaneous AFM topography and recognition imaging at the plasma membrane of mammalian cells, Seminars in Cell and Developmental Biology 73 (2018) 45 – 56.

[7] S. Cazaux, A. Sadoun, M. Biarnes-Pelicot, M. Martinez, S. Obeid, P. Bongrand, L. Limozin, P.-H. Puech, Synchronizing atomic force microscopy force mode and fluorescence microscopy in real time for immune cell stimulation and activation studies, Ultramicroscopy 160 (2016) 168–181.

[8] A. Miranda, M. Martins, P. A. A. De Beule, Simultaneous differential spinning disk fluorescence optical sectioning microscopy and nanomechanical mapping atomic force microscopy, Review of Scientific Instruments 86 (2015) 093705.

[9] S. Moreno Flores, J. L. Toca-Herrera, The new future of scanning probe microscopy: Combining atomic force microscopy with other surface-sensitive techniques, optical microscopy and fluorescence techniques, Nanoscale 1 (2009) 40–49.

[10] L. Zhou, M. Cai, T. Tong, H. Wang, Progress in the correlative atomic force microscopy and optical microscopy, Sensors 17 (2017) 938.

[11] R. Kassies, K. O. Van Der Werf, A. Lenferink, C. N. Hunter, J. D. Olsen, V. Subramaniam, C. Otto, Combined AFM and confocal fluorescence microscope for applications in bio-nanotechnology, Journal of Microscopy 217 (2005) 109–116.

[12] A. B. Mathur, G. A. Truskey, W. M. Reichert, Total internal reflection microscopy and atomic force microscopy (TIRFM-AFM) to study stress transduction mechanisms in endothelial cells, Critical Reviews in Biomedical Engineering 28 (2000) 197–202.

[13] D. Hu, M. Micic, N. Klymyshyn, Y. D. Suh, H. P. Lu, Correlated topographic and spectroscopic imaging beyond diraction limit by atomic force microscopy metallic tip-enhanced near-field fluorescence lifetime microscopy, Review of Scientic Instruments 74 (2003) 3347–3355.

[14] J. Yu, J. Yuan, X. Zhang, J. Liu, X. Fang, Nanoscale imaging with an integrated system combining stimulated emission depletion microscope and atomic force microscope, Chinese Science Bulletin 58 (2013) 4045–4050.

[15] J. V. Chacko, C. Canale, B. Harke, A. Diaspro, Sub-diffraction nano manipulation using STED AFM, PloS One 8 (2013) e66608.

[16] A. Monserrate, S. Casado, C. Flors, Correlative atomic force microscopy and localization-based super-resolution microscopy: Revealing labelling and image reconstruction artefacts, ChemPhysChem 15 (2014) 647–650.

[17] P. D. Odermatt, A. Shivanandan, H. Deschout, R. Jankele, A. P. Nievergelt, L. Feletti, M. W. Davidson, A. Radenovic, G. E. Fantner, High-resolution correlative microscopy: Bridging the gap between single molecule localization microscopy and atomic force microscopy, Nano Letters 15 (2015) 4896–4904.

[18] L. M. Hirvonen, S. Cox, STORM without enzymatic oxygen scavenging for correlative atomic force and fluorescence superresolution microscopy, Methods and Applications in Fluorescence 6 (2018) 045002.

[19] P. Bondia, R. Jurado, S. Casado, J. M. Domínguez-Vera, N. Gálvez, C. Flors, Hybrid nanoscopy of hybrid nanomaterials, Small 13 (2017) 1603784.

[20] M. G. L. Gustafsson, Surpassing the lateral resolution limit by a factor of two using structured illumination microscopy, Journal of Microscopy 198 (2000) 82–87.

[21] P. Kner, B. B. Chhun, E. R. Griffis, L. Winoto, M. G. L. Gustafsson, Super-resolution video microscopy of live cells by structured illumination, Nature Methods 6 (2009) 339.

[22] L. M. Hirvonen, K. Wicker, O. Mandula, R. Heintzmann, Structured illumination microscopy of – a living cell, European Biophysics Journal 38 (2009) 807–812.

[23] E. Betzig, Imaging life at high spatiotemporal resolution, in: Optics in the Life Sciences, Optical Society of America, 2015, p. JM1A.2.

[24] R. Förster, K. Wicker, W. Müller, A. Jost, R. Heintzmann, Motion artefact detection in structured illumination microscopy for live cell imaging, Opt. Express 24 (2016) 22121–22134.

[25] R. Förster, W. Müller, R. Richter, R. Heintzmann, Automated distinction of shearing and distortion artefacts in structured illumination microscopy, Optics Express 26 (2018) 20680–20694.

[26] L. Chopinet, C. Formosa, M. Rols, R. Duval, E. Dague, Imaging living cells surface and quantifying its properties at high resolution using AFM in QI™ mode, Micron 48 (2013) 26–33.

[27] L. Cong, F. A. Ran, D. Cox, S. Lin, R. Barretto, N. Habib, P. D. Hsu, X. Wu, W. Jiang, L. A. Marra ni, F. Zhang, Multiplex genome engineering using CRISPR/Cas systems, Science (New York, N.Y.) 339 (2013) 819–823.

[28] F. A. Ran, P. D. Hsu, J. Wright, V. Agarwala, D. A. Scott, F. Zhang, Genome engineering using the CRISPR-Cas9 system, Nature Protocols 8 (2013) 2281.

[29] D. Dambournet, S. H. Hong, A. Grassart, D. Drubin, Tagging endogenous loci for live-cell fluorescence imaging and molecule counting using ZFNs, TALENs, and cas9, Methods in Enzymology 546C (2014) 139–160.

[30] L. H. Schaefer, D. Schuster, J. Schaffer, Structured illumination microscopy: artefact analysis and reduction utilizing a parameter optimization approach, Journal of Microscopy 216 (2004) 165–174.

[31] R. Heintzmann, T. Huser, Super-resolution structured illumination microscopy, Chemical Reviews 117 (2017) 13890–13908.

[32] J. Demmerle, C. Innocent, A. J. North, G. Ball, M. Müller, E. Miron, A. Matsuda, I. M. Dobbie, Y. Markaki, L. Schermelleh, Strategic and practical guidelines for successful structured illumination microscopy, Nature Protocols 12 (2017) 988.

[33] C. Karras, M. Smedh, R. Förster, H. Deschout, J. Fernandez-Rodriguez, R. Heintzmann, Successful optimization of reconstruction parameters in structured illumination microscopy a practical guide, Optics Communications 436 (2019) 69 – 75.

[34] T. Ludwig, R. Kirmse, K. Poole, U. S. Schwarz, Probing cellular microenvironments and tissue remodeling by atomic force microscopy, Pflügers Archiv - European Journal of Physiology 456 (2008) 29–49.

[35] L. Novotny, B. Hecht, Principles of Nano-Optics, Cambridge University Press, 2006.

[36] P.-E. Mazeran, L. Odoni, J.-L. Loubet, Curvature radius analysis for scanning probe microscospy, Surface Science 585 (2005) 25–37.

[37] M. G. Gustafsson, L. Shao, P. M. Carlton, C. J. R. Wang, I. N. Golubovskaya, W. Z. Cande, D. A. Agard, J. W. Sedat, Three-dimensional resolution doubling in wide-field fluorescence microscopy by structured illumination, Biophysical Journal 94 (2008) 4957–4970.

[38] D. R. Stamov, S. B. Kaemmer, A. Hermsdörfer, J. Barner, T. Jähnke, H. Haschke, BioScience AFM – capturing dynamics from single molecules to living cells, Microscopy Today 23 (2015) 18–25.

